# Heli-SMACC: Helicase-targeting SMAll Molecule Compound Collection

**DOI:** 10.1101/2024.07.04.602122

**Authors:** Holli-Joi Martin, Mohammad A. Hossain, James Wellnitz, Enes Kelestemur, Joshua E. Hochuli, Sumera Parveen, Cheryl Arrowsmith, Timothy M. Willson, Eugene Muratov, Alexander Tropsha

## Abstract

Helicases have emerged as promising targets for the development of antiviral drugs; however, the family remains largely undrugged. To support the focused development of viral helicase inhibitors we identified, collected, and integrated all chemogenomics data for all available helicases from the ChEMBL database. After thoroughly curating and enriching the data with relevant annotations we have created a derivative database of helicase inhibitors which we dubbed Heli-SMACC (<u>Heli</u>case-targeting <u>SMA</u>ll Molecule <u>C</u>ompound <u>C</u>ollection). The current version of Heli-SMACC contains 20,432 bioactivity entries for viral, human, and bacterial helicases. We have selected 30 compounds with promising viral helicase activity and tested them in a SARS-CoV-2 NSP13 ATPase assay. Twelve compounds demonstrated ATPase inhibition and a consistent dose-response curve. The Heli-SMACC database may serve as a reference for virologists and medicinal chemists working on the development of novel helicase inhibitors. Heli-SMACC is publicly available at https://smacc.mml.unc.edu.

**Highlights:**

- We created a curated Helicase-Targeting SMAll Molecule Compound Collection (Heli-SMACC).
- Heli-SMACC covers 29 human, viral, and bacterial helicases.
- Twelve of thirty selected compounds demonstrated inhibitory activity in a SARS-CoV-2 NSP13 ATPase Assay.
- Heli-SMACC is freely available online at https://smacc.mml.unc.edu.

**TOC Graphic:** 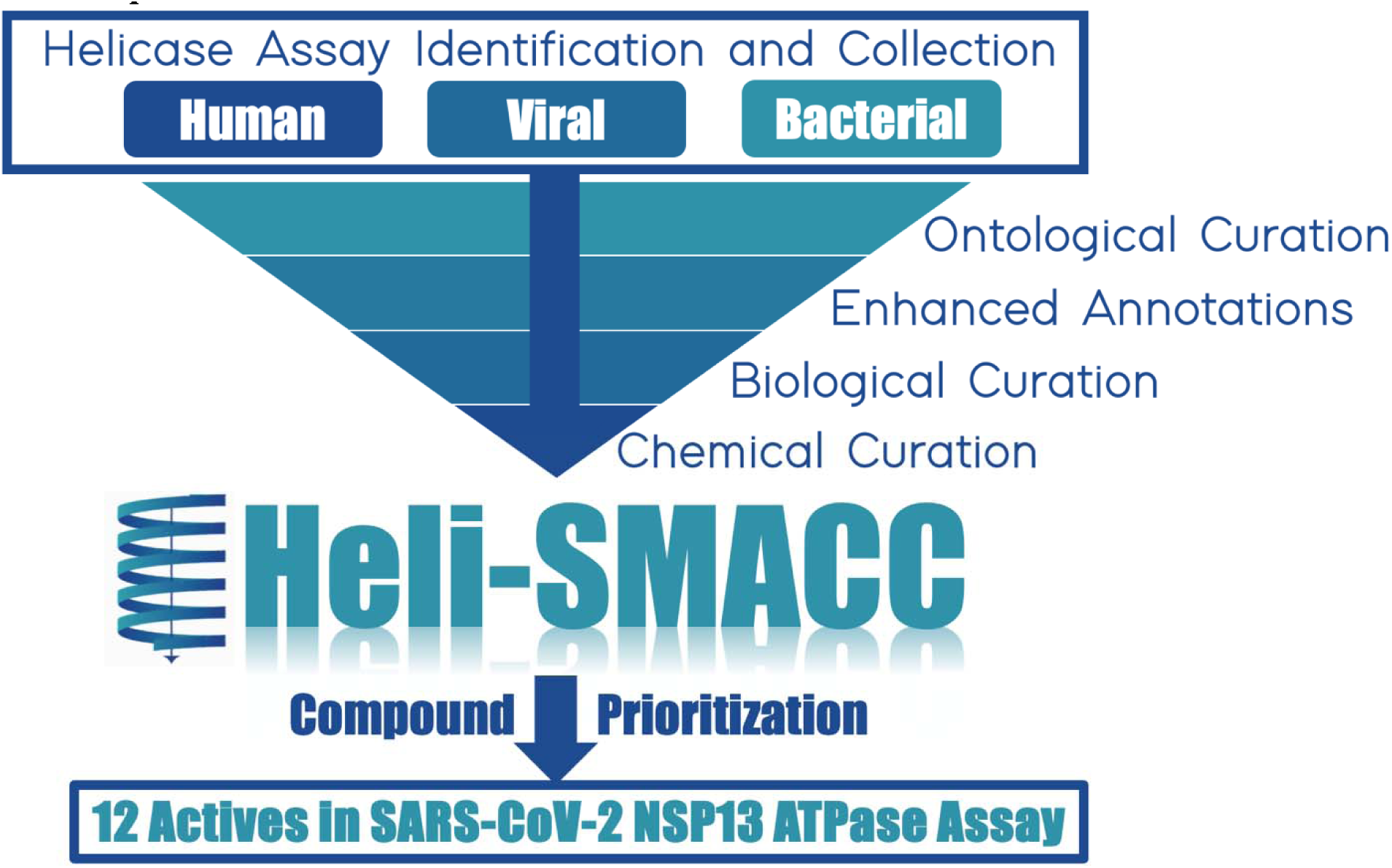

## 1. Introduction

DNA and RNA helicases are ubiquitous motor enzymes found in all domains of life from bacteria to eukaryotes. Helicases have emerged as promising targets for developing antiviral drugs, with the discovery that the replication of herpes viruses resistant to polymerase inhibitors, remain permissive to helicase inhibitors (Field and Biswas, 2011). However, while a few viral and human helicase inhibitors have entered clinical development, the family remains largely undrugged. All DNA and RNA helicases contain a common core structural fold (RecA), which serves as an ATP-binding domain. RecA contains specific motifs that have been used to classify helicases into six superfamilies (SF1-SF6) (Caruthers and McKay, 2002). All superfamilies contain motifs I and II (Walker A and Walker B), which are involved in the binding and hydrolysis of ATP, as well as an arginine finger (R), which aids in coupling the ATP hydrolysis reaction into a conformational change (Fairman-Williams et al., 2010). The two largest superfamilies are SF1 and SF2; helicases in these superfamilies share twelve out of thirteen motifs which can fold into RecA-like domains at the N- and C-terminals. Thus, regardless of the organism, helicases in the same superfamily will have some conserved structural features.

Although the cores of eucaryotic and viral helicases share a common RecA-like fold, the large difference in primary sequences increases the likelihood of specificity for viral over human helicases and low toxicity for the development of antiviral drugs (Kwong et al., 2005). However, in order to develop new antiviral helicase inhibitors, it is imperative to understand the full chemogenomic profiles of existing helicase inhibitors across all species. If possible, the identification of compounds with cross-helicase activity may distinguish key structural features of helicase inhibitors. This knowledge may aid in identifying hit chemotypes and in medicinal chemistry campaigns aiming to improve the antiviral selectivity of existing helicase inhibitors.

To support the development of antiviral helicase inhibitors, we collected and cleaned all publicly accessible data for compounds tested against any helicase and integrated it into a single highly curated, annotated, and publicly available database. We dubbed this database the Helicase-targeting SMAll molecule Antiviral Compound Collection (Heli-SMACC) and made it publicly available online at https://smacc.mml.unc.edu. We expect the Heli-SMACC database to support further computational and experimental medicinal chemistry studies targeting rational design and discovery of novel helicase inhibitors. To demonstrate this, we have selected 30 compounds with promising viral helicase activity and tested them in a SARS-CoV-2 NSP13 ATPase assay. Twelve compounds demonstrated ATPase inhibition and a consistent dose-response curve.

## 2. Methods

### 2.1 Data Curation

All data were extracted from ChEMBL 32, released on March 01, 2023 (Gaulton et al., 2017). The keyword “helicase” was used as the search query. A total of 269 helicase assays were identified, and all 21,076 associated bioactivities were downloaded. The data was pre-processed and curated following well-established protocols for chemical and biological data curation described elsewhere (Fourches et al., 2016, 2015, 2010). We transformed all possible bioactivity results into uM or % units and eliminated assays with irrelevant endpoints (i.e., K (/min) and Vmax (uM/s)). We set a binary threshold for all bioactivity results so that an assay result of 10 uM or >50% was considered an active result. Proper assay annotations were manually entered for entries that did not include the assay organism in the provided field (53.74% of entries), as well as for the assay target (53.44% of entries). We selected assays with the following activity standard types assigned by ChEMBL: Activity (30 entries), IC50 (1,220 entries), Potency (19,277 entries), and Inhibition (345 entries). We standardized the assay ontology for each assay target and organism, ensuring no two identical targets or organisms were represented differently in the database. We performed our standard workflow for chemical curation (Tropsha, 2010). Briefly, this included the removal of mixtures, inorganics, and salts, as well as the normalization of specific chemotypes. The resulting database (20,432 entries) was enhanced with additional ontologies related to the assay’s helicase including whether the helicase preferred DNA or RNA, and the helicase’s superfamily, subfamily, and function, where possible. Each assay result was kept in the database if the compound was tested in multiple assays (i.e., no duplicates were removed). This resulted in 5,976 entries labeled as active, 7,489 inactive, and 6,967 entries where the assay result was pre-labeled in ChEMBL as inconclusive. The summary of our database entries after curation is included in **Table 1**. While the inconclusive assay results remain in our database, only the actives and inactives are considered for the analyses in this manuscript.

**Table 1.** Summary of database entries after each step of data curation.

| **Curation Step** | **Number of Entries** |
| --- | --- |
| Data Extraction | 21,076 |
| Unit Transformation | 20,940 |
| Target and Organism Annotations | 20,934 |
| Assay Endpoint Selection | 20,872 |
| Removal of Mixtures, Inorganics, and Salts | 20,486 |
| Standardizing Chemotypes | 20,432 |

### 2.2 Helicase Inhibitor Selection

Here, we demonstrate one of many approaches to compound prioritization, in which we only consider active assay results in the analysis. To do so, we first isolated all 5,976 entries labeled as active after our data curation in section 2.1, which was a total of 4,081 unique compounds. To understand the broader activity profiles of these compounds, which may indicate the specificity/promiscuity of the compounds for other targets, we searched ChEMBL for all assays associated with each compound using the ChEMBL IDs as the search query. We followed the data curation procedure described above for the entire dataset. We stratified the assay results by compound and summarized the number and type of helicase and non-helicase targets against which the compounds were reported to be active. From this list, we created a subset of compounds with the objective of identifying promising viral helicase inhibitors. We used the STOPLIGHT hit scoring calculator (Wellnitz et al., 2023) to score and predict the important molecular properties and removed all compounds with a red STOPLIGHT score. To remove compounds with known promiscuity we filtered the list to include only compounds with five or fewer off-target activities reported. Finally, we filtered the compounds based on their availability on Molport. The final dataset is reported below in the Results section and Supplemental File 1.

## 3. Results

### 3.1 Overview of Heli-SMACC Data

There are assay results for 29 unique helicases in Heli-SMACC, spanning three broad organisms: humans, viruses, and bacteria (**Figure 1A**). While the viral helicases were the most represented in Heli-SMACC, the majority of assay results were for human helicases. In fact, over 91% of compounds present in Heli-SMACC were tested against one of nine human helicases. In contrast, compounds tested against viral or bacterial helicases account for only 6.1% and 1.7%, respectively, with ∼1% of compounds overlapping between any of the three organism classes (**Figure 1B**). Three of nine human helicases constitute 89% of the entire database (**Table S1)**: ATP-dependent DNA helicase Q1 (39.75%), Bloom syndrome protein helicase (30.30%), and Werner syndrome ATP-dependent helicase (18.90%). The human and bacterial helicase assay results were roughly split between actives and inactives (active constitute 46% and 53%, respectively), whereas only 23% of viral helicase assay results were active (**Figure 1C**).

**Figure 1.**
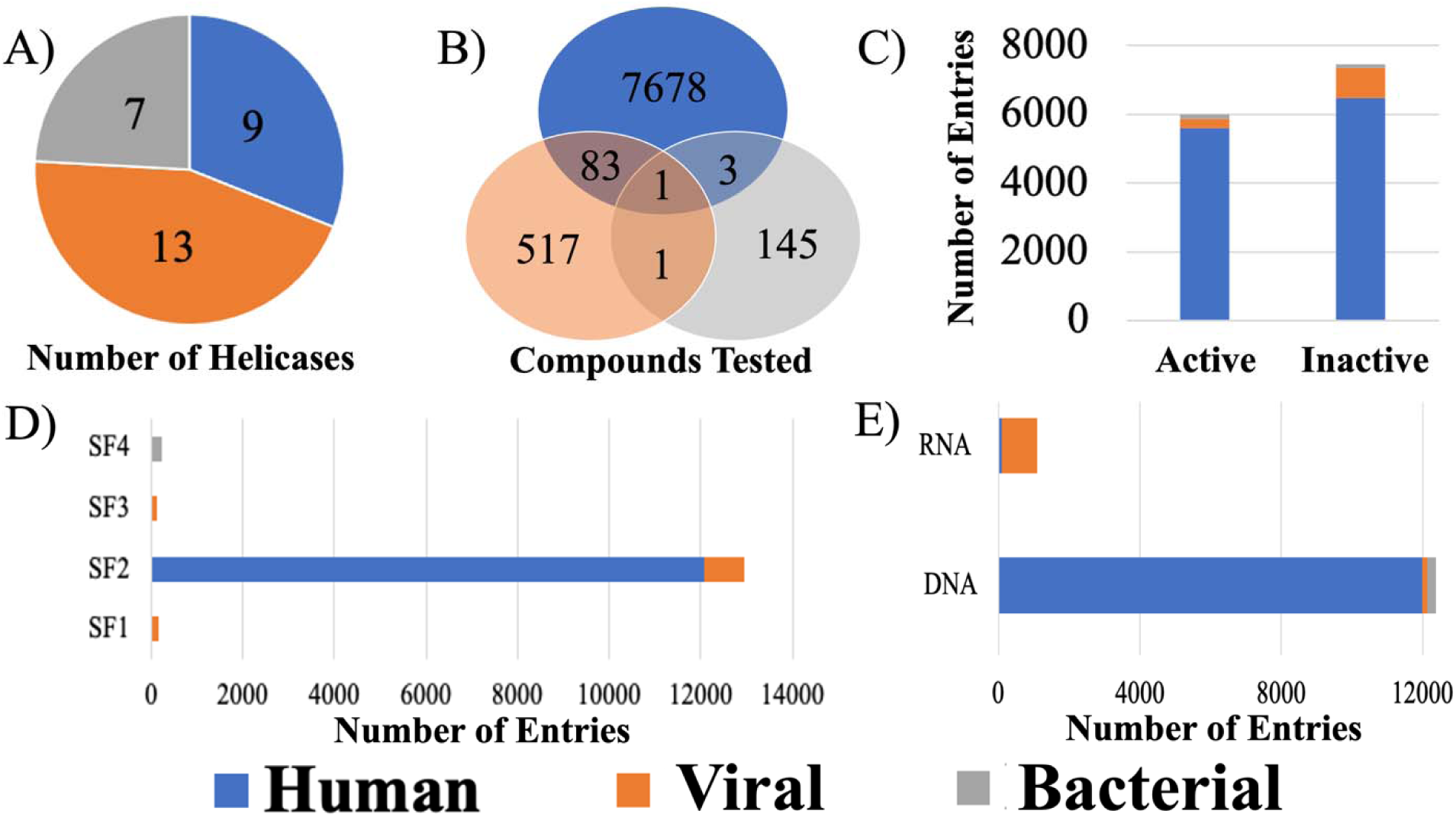
Summary of active and inactive human, viral, and bacterial helicase assay results in Heli-SMACC. A) Number of unique helicases present in each organism. B) Number of compounds tested across each helicase organism. C) Distribution of active and inactive assay results per organism. D) Number of entries present per helicase super family. E) Number of entries represented for DNA and RNA helicases.

Of the 1,162 assay entries associated with the 602 compounds targeting a viral helicase, ∼75% were associated with a helicase protein of a flavivirus, primarily hepatitis C virus (HCV) (783 entries), but also West Nile Virus (WNV) (51), Japanese Encephalitis Virus (JEV) (28), and Dengue Virus (DENV) (13). Also, ∼11% of entries were targeting the SARS-CoV-1 helicase protein. A minority of entries were associated with other viruses: Human papillomavirus type 11 (52), JC polyomavirus (35), BK polyomavirus (34), Human herpesvirus 1 (33), Human poliovirus 1 (1), and Enterovirus A71 (1).

The helicases present in Heli-SMACC represented four of six helicase superfamilies. Superfamily two (SF2) was the most represented and contained helicase data from both humans (12,065 entries) and viruses (875 entries). Superfamily one (SF1) also contained helicase data from both humans (3 entries) and viruses (164 entries). Whereas superfamily three (SF3) contained only 123 viral helicase entries, and superfamily four (SF4) contained only 234 bacterial helicase entries (**Figure 1D**). All of the bacterial helicase and 99.27% of the human helicase entries had a specificity for DNA (**Figure 1E**). Conversely, 86.7% of the viral helicase entries were for helicases with a specificity for RNA.

### 3.2 Analysis of Compounds in Heli-SMACC

Compounds associated with active and inactive helicase assay results were mapped in chemical space and grouped by the helicase assay organism: human, viral, or bacterial (**Figure 2A**). All compound similarity maps were generated using OSIRIS DataWarrior software (Sander et al., 2015). Given the over-representation of human helicase data in Heli-SMACC, the majority of compound data points in chemical space are green. However, the three major organism classes (human, viral, bacterial) formed distinct clusters in chemical space. The largest clusters from each major organism class are circled in their relative colors (human (green), viral (orange), and bacterial (blue)). Representative molecules of each cluster are shown in **Figure 2B**. For the human helicases, cluster 1 consists of pyrimidine-dione molecules, and cluster 2 represents the polyphenol class of molecules. From our analysis of the representative molecules, cluster 1 is more likely to inhibit helicases, whereas cluster 2 appears to be non-specific, given that polycyclic polyphenolic compounds are not druglike molecules and are often identified as false positives. The two major compound clusters from the viral helicase inhibitors were a series of hydrazines (cluster 1) and triazoles (cluster 2). Visual inspection of viral helicase cluster 1 revealed several potential helicase inhibitors with druglike properties. Finally, the compounds targeting bacterial helicases are primarily of the chromen (cluster 1) or pyrazine (cluster 2) templates. Compared to chromens, pyrazines are more druglike molecules with lower molecular weight and higher potential for selectivity. In summary, we found that the most represented chemical motifs differed between each organism class.

**Figure 2.**
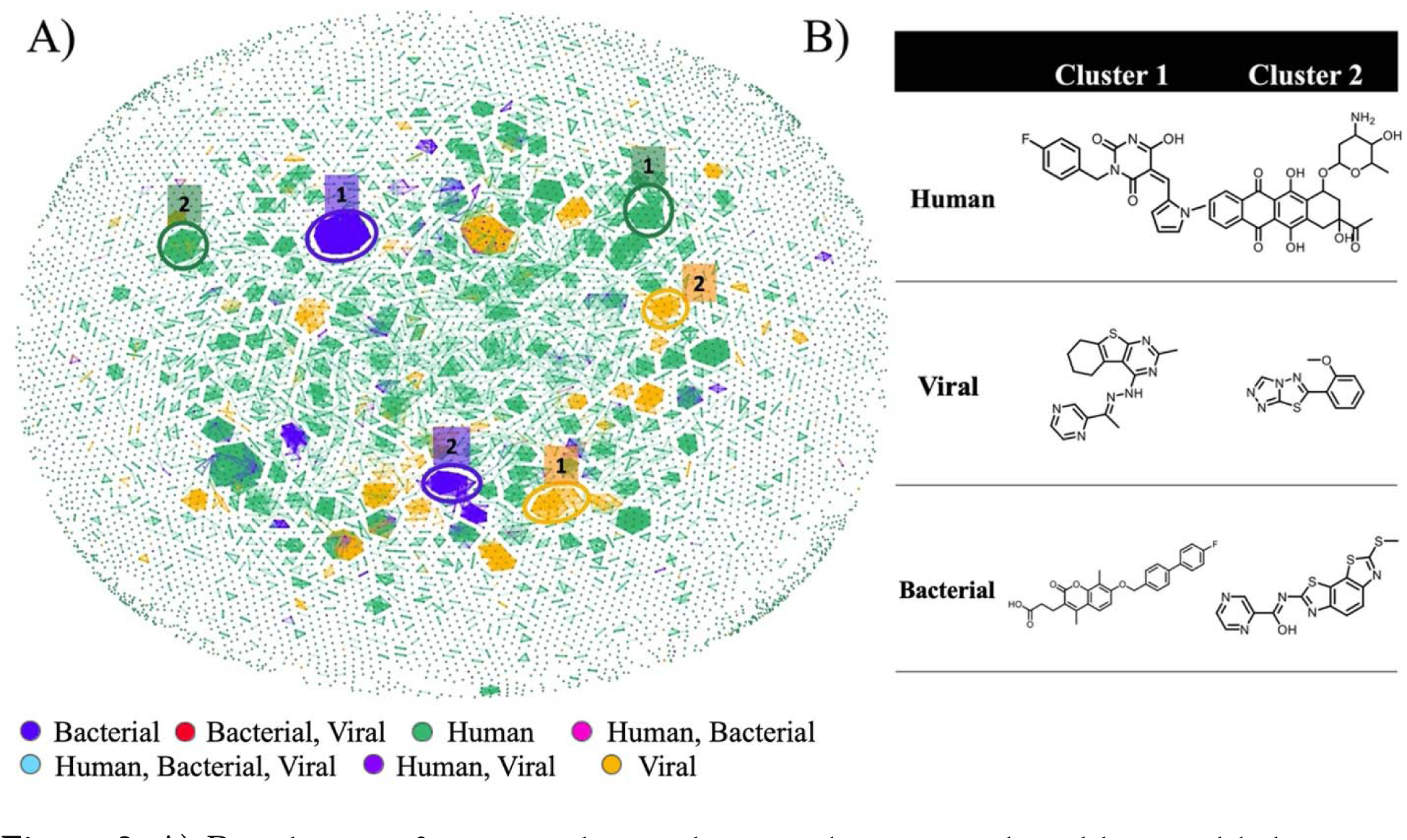
A) Distribution of compounds tested against human, viral, and bacterial helicases in chemical space. The two major clusters are highlighted, corresponding to the data color for the human (green), viral (orange), and bacterial (blue) helicase data. B) Representative molecules from the top clusters of compounds tested against human, viral, and bacterial helicases.

The number of compounds tested against helicases in SF1–SF4 is shown in **Figure 3A**. It is clear that the majority of compounds in Heli-SMACC are tested against SF2; however, there were 67 compounds tested against multiple helicase superfamilies. We looked at the chemical space of each helicase superfamily (**Figure 3B**) and saw that each one formed a distinct cluster with very few overlapping clusters from different superfamilies. The top clusters are circled in **Figure 3B** and colored relative to each superfamily: SF1 (dark blue), SF2 (red), SF3 (purple), and SF4 (light green). A representative molecule from each top cluster is shown (**Figure 3C**). Cluster 1 of the SF1 superfamily contains a very distinct class of molecules identified predominantly by the halogenated octahydro-purin moiety. These compounds are derivatives of purine analogs and potential hits for several disease targets, including helicases. The most represented molecules from SF2 (cluster 2) were pyrimidine and hydroxypyrimidines; these are drug-like molecules that have been frequently reported to act at multiple human and viral targets. The SF3 superfamily contains a cluster of substituted triazolo-thiadiazole class of molecules (cluster 3). Finally, SF4 is represented by cluster 4 and contains a chromen core with substituted propanoic acid derivatives.

**Figure 3.**
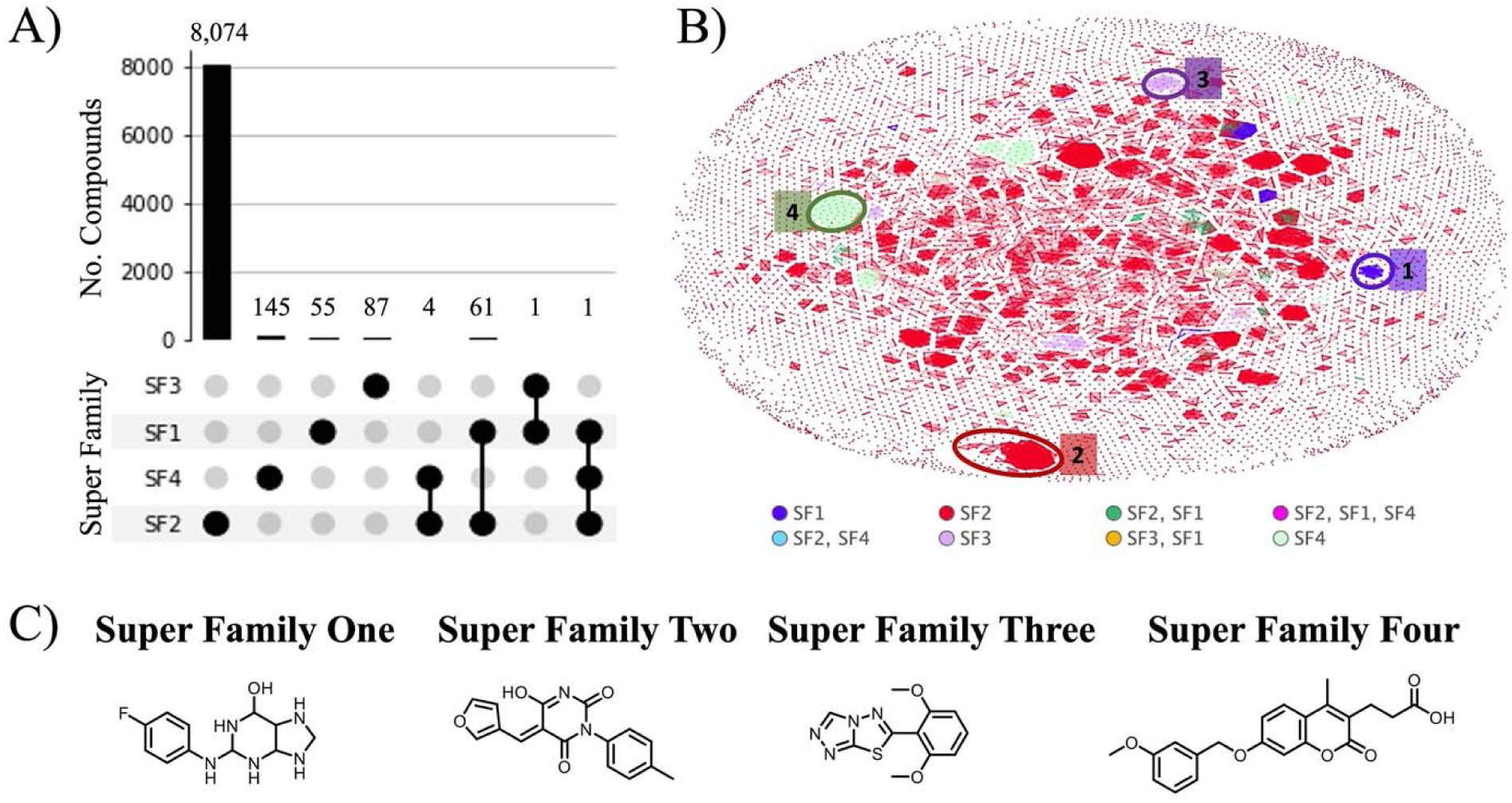
A) The number of compounds tested against helicases in superfamilies one through four. B) Distribution of compounds tested against each helicase superfamily in chemical space. The top clusters are highlighted in the corresponding data color for superfamilies one (dark blue), two (red), three (purple), and four (light green). C) Representative molecules from the top clusters of compounds tested super families one through four.

We also looked at the distribution of active compounds within chemical space (**Figure 4A**) using the OSIRIS DataWarrior software (Sander et al., 2015). The coloring scheme was limited to compounds with Tanimoto similarity greater than 0.8 to its nearest neighbor. Chemical similarity with a nearest neighbor less than 0.8 is indicated in grey, while a similarity score of 0.8 is indicated in red and 0.99 in lime green. The five major clusters of highly similar compounds are circled in black with a representative molecule from each of the top five clusters shown in **Figure 4B**. The top five clusters represent compounds with the core of pyrimidine (**Figure 4B, 1**), benzothiazoles (**Figure 4B, 2**), anthracycline (**Figure 4B, 3**), chromen (**Figure 4B, 4**), and polyphenolic chromen (**Figure 4B, 5**).

**Figure 4.**
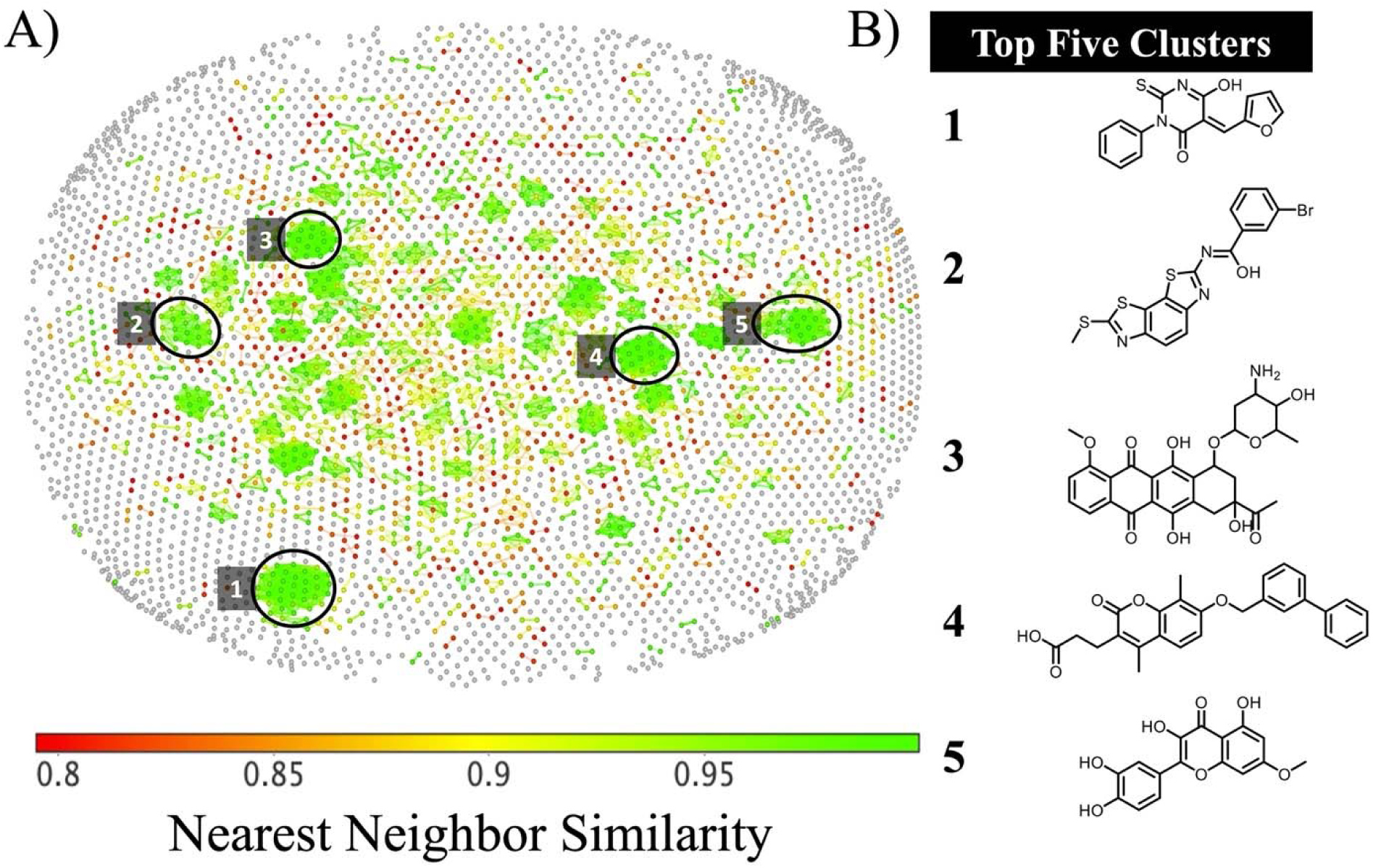
A) Distribution of compounds with reported helicase activity in chemical space. Grey coloring indicated chemical similarity to nearest neighbor <0.8, whereas a coloring scale from red to green indicated >0.8 similarity between nearest neighbors. The five major clusters of highly similar compounds are highlighted in black. B) Representative molecules from the top five clusters.

### 3.3 Identification of Promising Helicase Inhibitors

With a goal of identifying helicase inhibitor chemotypes with activity against viruses with high pandemic potential, 177 compounds from Heli-SMACC exemplifying the viral helicase inhibitors were flagged for experimental testing (**File S1 Tab 1**). Initially, a subset of these inhibitors was purchased from Molport and tested in a SARS-CoV-2 NSP13 ATPase assay (**see Supplemental Methods**).

#### 3.3.1 Viral Helicase Inhibitors

Of the 177 commercially available compounds that were flagged for experimental testing (examples in **Figure 5**), 101 were inhibitors of hepatitis C virus (HCV) helicase, including 5 compounds that inhibited HCV plus one or more flavi- or corona-virus helicase(s). We also selected 30 human papillomavirus (HPV) inhibitors, 11 SARS-CoV-1 inhibitors, 11 herpes simplex virus (HSV) inhibitors, nine flavivirus inhibitors (WNV, JEV, DENV) and a handful of compounds inhibiting other viruses. The selected chemotypes consist of natural product derivatives to drug-like small molecules. We failed to find a correlation between the chemical nature of the identified hits and its viral superfamily (**Figure 5**). Therefore, to map the selectivity and potency across viral helicases, we intend to screen them against a panel of viruses and in biochemical assays (**Table 2**).

**Figure 5.**
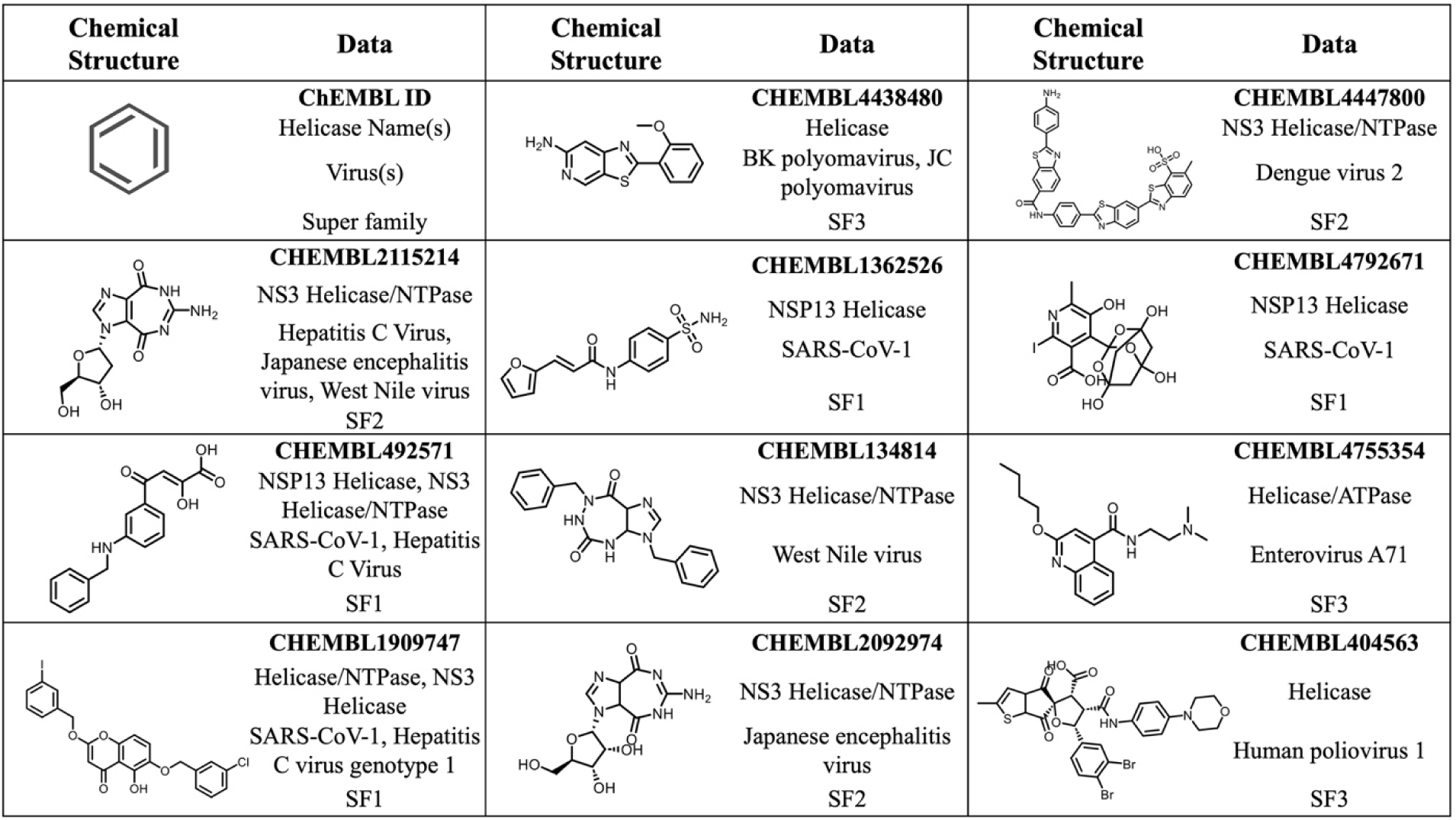
Examples of viral RNA helicase inhibitors nominated for experimental testing. The upper left cell defines the tables contents.

**Table 2.** Plan of experimental assays for the selected helicase inhibitors in super family 1 helicases.

| Superfamily | Virus | Biochemical assays | In-vitro |
| --- | --- | --- | --- |
| SF1 | SARS-CoV-2 | NSP13 ATPase,<br>Unwindase, SPR | Cell-based assay |
| SF1 | MERS | SPR |  |
| SF1 | CHIKV | NSP2 NTPase | Cell-based assay |

#### 3.3.2 Biological Activity of Experimentally Tested Helicase Inhibitors

For our first round of testing, we purchased 30 out of 177 compounds that we prioritized based on their reported viral helicase inhibition. These compounds were screened in a SARS-CoV-2 NSP13 ATPase assay at 50 µM concentration. A subset of 28 compounds inhibited ATPase by more than 40%. The subsequent dose-dependent assay of these 28 compounds showed an IC_50_ against the NSP13 ATPase ranging from 3 to 77 µM (**Figure 6**). As evident from **Figure S1**, several compounds showed scattered dose-responses due to their poor aqueous solubility profile (**File S1 Tab 2**). These compounds will also be tested for their aggregation in the assay media.

**Figure 6.**
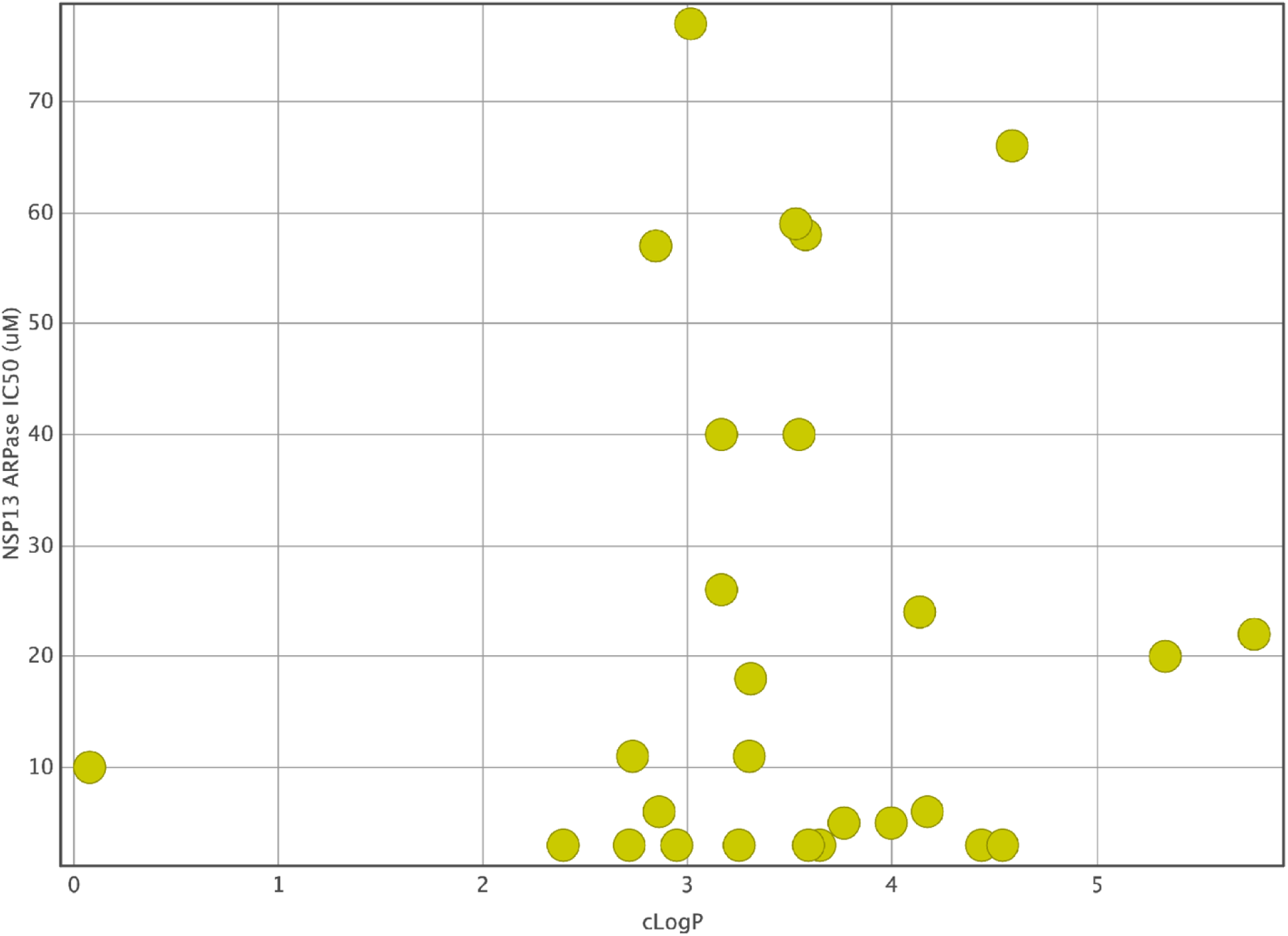
Dose-response of selected helicase inhibitors in NSP13 ATPase assay (kinase-glo). The Y-axis shows the % Activity, and the X-axis shows the concentration of each compound.

## 4. Discussion

By collecting, curating, and integrating all publicly accessible data for compounds targeting helicases, we have used the data available in ChEMBL, an online collection of bioactive molecules with drug-like properties (Gaulton et al., 2017), to build a curated, annotated, and publicly available database of compounds tested against any helicase, called the Helicase-targeting SMAll molecule Antiviral Compound Collection (Heli-SMACC) and made it publicly available online at https://smacc.mml.unc.edu. This work is an extension to our SMACC database (Small Molecule Antiviral Compound Collection) which contains over 32,500 entries for 13 viruses of high pandemic potential, tested in both phenotypic and target-based assays (Martin et al., 2023). By publishing SMACC, we aimed to increase the clarity and accessibility of antiviral assays and the relevant, associated information to a broader scientific community. The Heli-SMACC database was created in that same regard and is a rich source of compound bioactivity data associated with helicase assays.

Despite the presence of helicase proteins in every living organism, only 29 helicases have experimental data reported in the ChEMBL database, with 48% of those helicases having fewer than 70 data points available. This scarcity of data is likely attributable to the challenges associated with developing and validating assays that accurately measure helicase activity. Traditionally, the ATPase assay has been the primary method employed to monitor the inhibition of helicases by small molecules. This assay works by measuring the conversion of ATP to ADP, a reaction essential for the function of helicases. Another commonly used method is the helicase unwindase assay, which directly assesses the potency of helicase inhibitors in an *in-vitro* setting. Biophysical assays such as Surface Plasmon Resonance (SPR) have recently emerged. However, these novel methods have been plagued by a lack of reproducibility, which we have experienced first-hand in our drug discovery campaigns. When using all the assays concurrently, we have observed inconsistent results, pointing towards a multifaceted problem with isolated, truncated, and engineered proteins, as helicase fragments do not exhibit the same behavior in isolation as they do in their natural cellular environment. This discrepancy underscores the need for further optimization of bioassays to replicate the data more accurately. Moreover, it is of paramount importance to correlate helicase activities with the *in-vitro* antiviral potency of each identified hit.

As efforts are made to improve the reproducibility of compound bioactivities in helicase assays and new data emerges, we will continue to update the Heli-SMACC database, ensuring consistency in data curation, annotation, and proper ontological harmonization. In addition to updated content, the SMACC and Heli-SMACC databases will undergo a transformation in their web interface this year to enable better searchability, filtering, and data exportation. Some planned updates include the ability to search and filter by chemical, chemical similarity, assay, organism, and more. Our update will also enable online search and filtering criteria to allow users the ability to isolate and export relevant subsets of the data. We expect these updates to greatly improve the user experience and provide quicker, more tailored access to the Heli-SMACC database.

In this manuscript, we demonstrate one approach for selecting promising viral helicase inhibitors from Heli-SMACC. However, we emphasize that there are many other approaches for querying Heli-SMACC depending on the user’s goals. We prioritized 177 compounds from Heli-SMACC for experimental testing (**File S1 Tab 1**). Initially, a subset of 30 viral helicase inhibitors was purchased from Molport and tested in a SARS-CoV-2 NSP13 ATPase assay. Most viral helicase inhibitors we selected for experimental testing were HCV inhibitors; unsurprisingly, due to their high representation in Heli-SMACC (∼2/3 of all viral helicase entries).

It is likely that a high sequence conservation exists across non-structural proteins required for viral replication within a viral clade, which has been demonstrated for the coronaviruses (Melo-Filho et al., 2022). We hypothesize that the HCV helicase inhibitors might also inhibit other flaviviruses such as Dengue, West Nile, and Yellow Fever viruses, mainly because of the conserved structural motifs between helicases within SF2. As mentioned above, helicases in SF1 and SF2 share 12 of 13 structural motifs. Furthermore, we wanted to determine if these inhibitors showed activity at SF1, which influenced our initial selection of compounds tested in the SARS-CoV-2 NSP13 ATPase assay. Interestingly, 93.33% of the compounds we selected showed NSP13 ATPase inhibition >40% at 50 µM, with the subsequent dose-dependent assay of 28 compounds showing IC_50_ ranging from 3 to 77 µM. From these, 12 compounds showed a consistent dose-response curve (**Figure S1**; **Table S2**), while the rest suffered from solubility issues (**File S1 Tab 2**). Additional cell-based screening and physicochemical property validation for the selected hits are in process, along with testing their ability to aggregate in the assay media.

Our initial experiment suggested that viral helicase inhibitors may show activity at both SF1 and SF2 helicases. We nominated 151 inhibitors (**File S1 Tab 3**) and tested 24 inhibitors of human SF2 helicases in the SARS-CoV-2 NSP13 ATPase assay to determine if they might also inhibit a viral SF1 helicase. Among these 41 compounds, 8 showed weak activity (29 to 90 µM) in the NSP13 ATPase assay (**File S1 Tab 4; Table S4**). This suggests viral helicases are more homologous to each other than to human helicases, which may aid in future efforts to design viral-specific helicase inhibitors, avoiding their human anti-targets.

## 5. Conclusions

To support the development of antiviral helicase inhibitors, we developed the Heli-SMACC (<u>Heli</u>case-targeting <u>SMA</u>ll molecule <u>C</u>ompound <u>C</u>ollection) database. This database contains 20,432 entries of bioactivity data for compounds targeting viral, human, and bacterial helicases. We prioritized 177 viral helicase inhibitors for experimental testing against viruses of pandemic potential. We initially tested 30 viral helicase inhibitors in a SARS-CoV-2 NSP13 ATPase assay. Interestingly, 93.33% of the compounds we selected showed activity greater than 40%, with the subsequent dose-dependent assay showing IC_50_ ranging from 3 to 77 µM. From these, 12 compounds showed consistent dose-response curves. The Heli-SMACC database built in this study may serve as a reference for virologists and medicinal chemists working to discover and develop novel helicase inhibitors. The Heli-SMACC database is publicly available in the form of a searchable Excel spreadsheet at https://smacc.mml.unc.edu.

## 7. Acknowledgments

Authors from UNC-Chapel Hill were supported by the National Institutes of Health (Grants U19AI171292 and R01GM140154). JW was supported by the National Institute of General Medical Sciences (Award T32GM135122). The content is solely the responsibility of the authors and does not necessarily represent the official views of the National Institutes of Health. HJM acknowledges the support from the American Foundation for Pharmaceutical Education’s (AFPE) Predoctoral Fellowship. The Structural Genomics Consortium is a registered charity (no: 1097737).

## 8. Data Availability

The curated Heli-SMACC database can be found at https://smacc.mml.unc.edu.

## Supporting Information

**Table S1.** Summary of all helicases present in Heli-SMACC.

| Helicase Name | Organism(s) | Super Family | DNA/RNA | Entries |
| --- | --- | --- | --- | --- |
| DNA2 Helicase/Nuclease | Homo sapiens | SF1 | DNA | 3 |
| DNA Helicase/Primase | Human herpesvirus 1, Herpes simplex virus type 1 | SF1 | DNA | 33 |
| Helicase/NTPase | SARS-CoV-1 | SF1 | RNA | 20 |
| NSP13 Helicase | SARS-CoV-1 | SF1 | RNA | 111 |
| NS3 Helicase/NTPase | Hepatitis C, Dengue Virus, West Nile virus, Japanese encephalitis virus | SF2 | RNA | 875 |
| ATP-dependent DNA helicase Q1 | Homo sapiens | SF2 | DNA | 5352 |
| ATP-dependent RNA helicase DDX1 | Homo sapiens | SF2 | RNA | 11 |
| ATP-dependent RNA helicase DDX3X | Homo sapiens | SF2 | RNA | 28 |
| BRR2 Helicase | Homo sapiens | SF2 | RNA | 6 |
| Bloom syndrome protein helicase | Homo sapiens | SF2 | DNA | 4080 |
| SUV3 Helicase | Homo sapiens | SF2 | RNA | 40 |
| Werner syndrome ATP-dependent helicase | Homo sapiens | SF2 | DNA | 2546 |
| eIF4A3 helicase | Homo sapiens | SF2 | RNA | 2 |
| Helicase | BK polyomavirus, Human poliovirus 1, JC polyomavirus | SF3 | DNA, RNA | 70 |
| Helicase/ATPase | Enterovirus A71 | SF3 | RNA | 1 |
| E1 DNA Helicase/ATPase | Human papillomavirus type 11 | SF3 | DNA | 52 |
| DNA Helicase | Bacillus anthracis, Staphylococcus aureus | SF4 | DNA | 158 |
| Helicase IV | Escherichia coli | SF4 | DNA | 6 |
| DNAc Helicase | Staphylococcus aureus | SF4 | DNA | 18 |
| DNAb Helicase | Staphylococcus aureus, Escherichia coli, Pseudomonas aeruginosa | SF4 | DNA | 52 |

**Figure S1.**
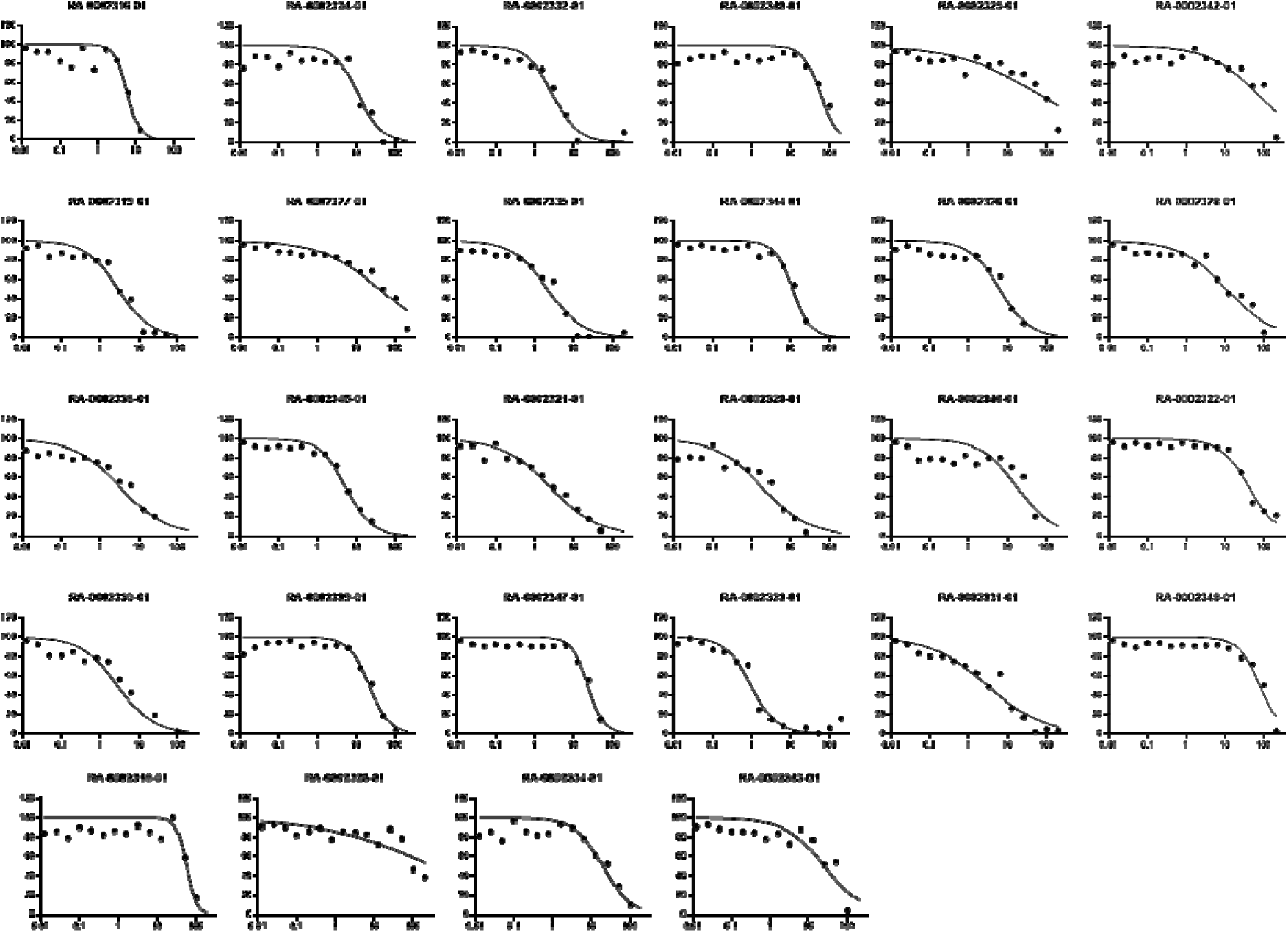
Dose-response of selected viral helicase inhibitors in NSP13 ATPase assay (kinase-glo).

**Table S2:**
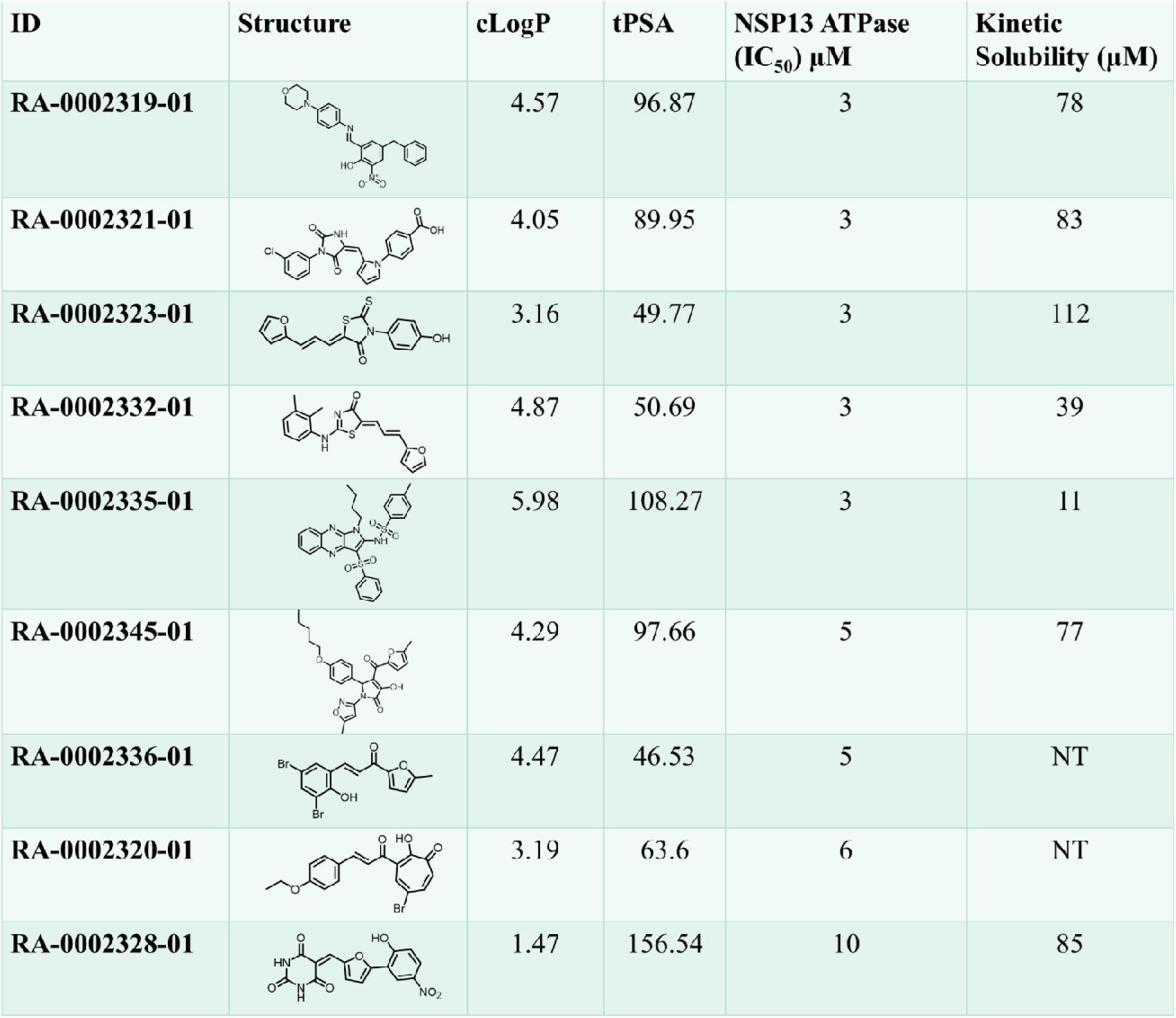
Calculated physicochemical properties and experimental kinetic solubility of the selected NSP13 ATPase assay hits from the viral helicase inhibitors.

| ID | Structure | cLogP | tPSA | NSP13 ATPase (IC<sub>50</sub>) μM | Kinetic Solubility (μM) |
| --- | --- | --- | --- | --- | --- |
| RA-0002319-01 |  | 4.57 | 96.87 | 3 | 78 |
| RA-0002321-01 |  | 4.05 | 89.95 | 3 | 83 |
| RA-0002323-01 |  | 3.16 | 49.77 | 3 | 112 |
| RA-0002332-01 |  | 4.87 | 50.69 | 3 | 39 |
| RA-0002335-01 |  | 5.98 | 108.27 | 3 | 11 |
| RA-0002345-01 |  | 4.29 | 97.66 | 5 | 77 |
| RA-0002336-01 |  | 4.47 | 46.53 | 5 | NT |
| RA-0002320-01 |  | 3.19 | 63.6 | 6 | NT |
| RA-0002328-01 |  | 1.47 | 156.54 | 10 | 85 |

### SARS-CoV-2 NSP13 ATPase assay protocol

The final reaction mixtures consisted of 50 mM HEPES, pH 7.5, 5% Glycerol, 5 mM magnesium acetate, 5 mM DTT, and 0.01% BSA, 0.1 nM nsp13, 3.5 nM ssDNA and 2.5 µM ATP. The reactions were started by the addition of substrates and incubated for 60 min at room temperature. The level of enzyme activity was then measured using a luciferase reagent, and the data were analyzed using GraphPad Prism 9.

### Solubility assay protocol

Kinetic solubility was performed by the CRO, Analiza. The reported methods for sample preparation and analysis as follows:

#### Sample Preparation

50-fold dilutions of each DMSO stock solution were prepared in singleton by combining 6μL of DMSO stock with 294μL of the appropriate media in a Millipore solubility filter plate with 0.45μM polycarbonate filter membrane using Hamilton Starlet liquid handling. The final DMSO concentration is 2.0% and maximum theoretical compound concentration is 200μM (assuming stock concentration of 10mM). The filter plate was heat sealed for the duration of the 24-hour incubation period.

#### Buffer Preparation

1XPBS, pH 7.4: Phosphate Buffered Saline solution 10X, PBS (Fisher Bioreagent part number BP399-500). 50mL was added to approximately 450mL HPLC grade H2O. The volume of the solution was then adjusted to 500mL for a total dilution factor of 1:10 and a final PBS concentration of 1X.

#### Solubility Analysis

The samples were placed on a rotary shaker (200RPM) for 24 hours at ambient temperature (20.3–22.3°C) then vacuum filtered. All filtrates were injected into the nitrogen detector for quantification on Analiza’s Automated Discovery Workstation. The results are reported here in μg/ml and μM.

#### Calculation of Results

The equimolar nitrogen response of the detector is calibrated using standards which span the dynamic range of the instrument from 0.08 to 4500 μg/ml nitrogen. The filtrates were quantified with respect to this calibration curve. The calculated solubility values are corrected for background nitrogen present in Analiza’s in-house DMSO and the media used to prepare the samples. Three separate on-board performance indicating standards were assayed in triplicate from 10mM stock solutions at 2.0% DMSO with the University of North Carolina supplied compounds, and all results were within the acceptable range. A comments field contains notes pertinent to the assay of each compound, such as below LOQ or measured solubility is greater than 75% of the dose concentration, the actual solubility may be higher.

### Multi-helicase Inhibitors

There were 151 compounds selected which demonstrated activity at multiple helicases (examples in **Figure S2**). The majority of these multi-helicase inhibitors targeted a combination of Bloom syndrome protein helicase, Werner syndrome ATP-dependent helicase, and ATP-dependent DNA helicase Q (examples in **Figure S2**). There was also a subset of compounds that showed activity at the viral Hepatitis C NS3 Helicase/NTPase as well as one of the human helicases (Bloom syndrome protein helicase, Werner syndrome ATP-dependent helicase, or ATP-dependent DNA helicase Q1). These pan helicase inhibitors are intriguing as a tool molecule to investigate the role of helicase and underscore the homology of helicase across the species. All the helicases contain RecA type domain, and there is possibility of similar modes of action of ligands.

**Figure S2.**
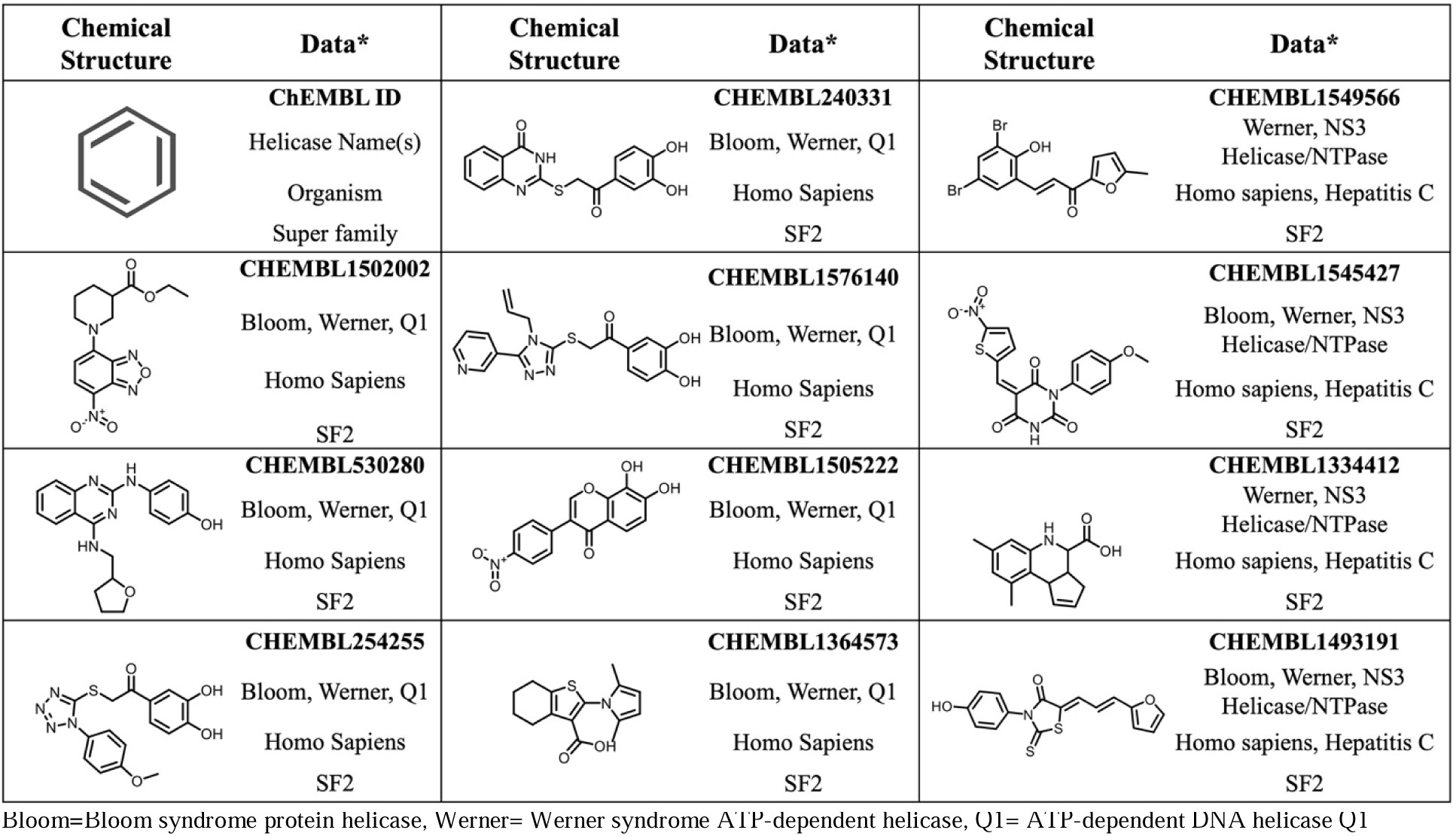
Examples of compounds active at two or more helicases nominated for experimental testing.

**Table S3:**
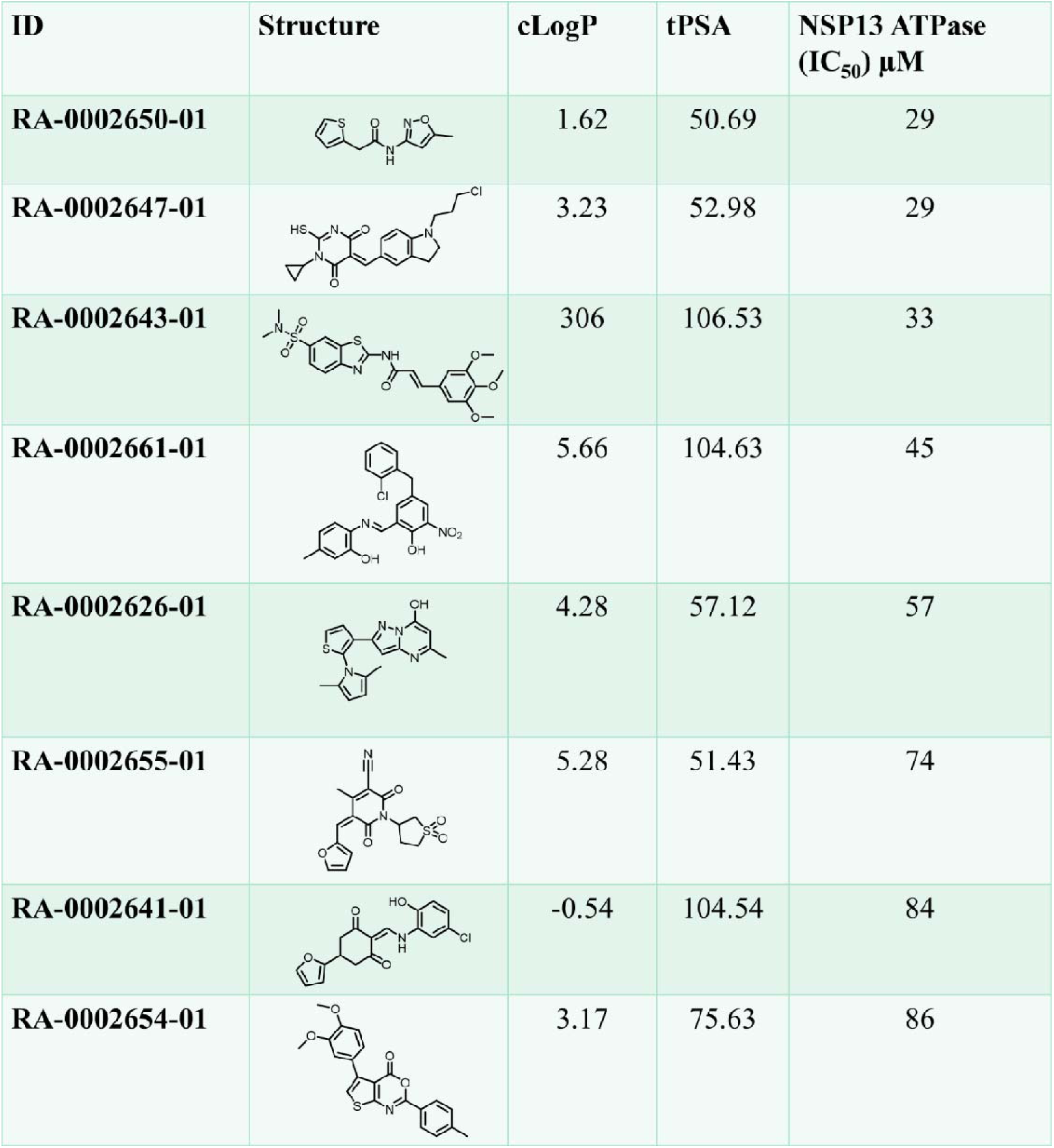
SARS-CoV-2. NSP13 ATPase inhibition of selected SF2 series compounds.

| ID | Structure | cLogP | tPSA | NSP13 ATPase (IC<sub>50</sub>) μM |
| --- | --- | --- | --- | --- |
| RA-0002650-01 |  | 1.62 | 50.69 | 29 |
| RA-0002647-01 |  | 3.23 | 52.98 | 29 |
| RA-0002643-01 |  | 306 | 106.53 | 33 |
| RA-0002661-01 |  | 5.66 | 104.63 | 45 |
| RA-0002626-01 |  | 4.28 | 57.12 | 57 |
| RA-0002655-01 |  | 5.28 | 51.43 | 74 |
| RA-0002641-01 |  | -0.54 | 104.54 | 84 |
| RA-0002654-01 |  | 3.17 | 75.63 | 86 |

